# The impact of microplastic contamination in cow manure on reproductive behavior and larval survival in the dung beetle *Onthophagus taurus*

**DOI:** 10.1101/2025.10.14.682200

**Authors:** Nathan McConnell, Jaydon Galindo Lovell, Jill Walker, Benjamin J. Mathews, Scott G. Morton, Jonathan B. Shurin, Patrick T. Rohner

## Abstract

Microplastics are an emerging environmental hazard on a global scale. Their detection in agricultural environments is of particular concern not only for food contamination, but also because microplastics negatively impact detritivores and their ecosystem functioning. Dung beetles in particular provide vital ecosystem services in agricultural environments and are often vulnerable to anthropogenic hazards, but whether they are affected by microplastics remains unclear. Here, we test whether artificial contamination of cow dung with thermoplastic polyurethane (TPU) has the potential to affect the juvenile development and maternal behavior of the bull-headed dung beetle *Onthophagus taurus*. Dung beetles exhibited high mortality when exposed to elevated concentrations of TPU. In addition, females were equally likely to provision offspring with TPU-spiked (and lethal) cow dung as with control dung, suggesting that females cannot differentiate between highly toxic microplastic-contaminated and uncontaminated cow dung. Our findings highlight potentially severe consequences for dung beetles if microplastics persist and accumulate, although the levels of exposure in the field are unknown. Although the direct environmental hazards and the mechanisms mediating the negative impacts of TPU microplastics remain to be assessed, this study suggests that microplastics may negatively impact dung beetles and their ecosystem services. Future work assessing exposure levels in the field as well as dung beetles’ potential to evolve resistance against microplastic pollution will be necessary to assess the long-term impact of microplastic presence on dung beetle ecosystem functioning.

## 1. Introduction

Invertebrate detritivores play a major role in ecosystem functioning in agricultural landscapes across the globe (Anderson et al. 2024; Nichols et al. 2008). Hundreds of species of beetles, flies, and other invertebrates contribute to nutrient cycling, soil health, plant growth, and the suppression of parasites (Skidmore 1991). However, their diversity, abundance, and ecological functions are increasingly threatened by anthropogenic pollutants, including veterinary antiparasitic drugs and insecticides (Floate et al. 2005; Kavanaugh and Manning 2020; Puniamoorthy et al. 2014) and other potentially harmful, more understudied contaminants, such as microplastics (MPs). Across terrestrial and aquatic ecosystems, various types of microplastics have been shown to harm invertebrate detritivores and disrupt their ecological functions. For instance, in soils, tire‐derived particles reduced the rate at which darkling beetles fragment organic matter, slowing down microbial decomposition (Staczek and Mbora 2025). Earthworms exposed to microplastic‐ spiked soils exhibit reduced growth and cocoon production suggesting non‐selective ingestion can drive bioaccumulation (Trakić et al. 2024). Although bacteria and fungi perform the bulk of soil organic-matter decomposition, they depend on detritivores, such as darkling beetles and earthworms, to fragment and moisten plant residues, aerate soils via burrowing, and disperse microbial inocula, thereby enabling efficient microbial breakdown (Culliney 2013; Jones et al. 1994; Lavelle et al. 1995). Microplastics have also been shown to slow leaf-litter decomposition in streams, thereby impairing organic-matter breakdown and nutrient release (Lopez-Rojo et al. 2020). Collectively, these examples demonstrate that microplastic contamination can impair detritivore functions, thereby disrupting microbial decomposition and undermining nutrient‐cycling.

Dung beetles (Scarabaeidae: Scarabaeinae) comprise a large group of scarabs (>6,000 species) that are common in all terrestrial ecosystems from tropical rain forests to deserts and alpine grasslands (Hanski and Cambefort 1991; Philips 2011). The presence of dung beetles improves pasture health because many species tunnel underneath vertebrate dung and construct underground brood chambers (often referred to as ‘*brood balls*’) out of dung, which serve as the ontogenetic environment for a single larva. Through the excavation of breeding tunnels and the relocation of dung underground (tunnels are often 20-30 cm deep but can be more than 1 meter in depth (Kingston and Coe 2009)), dung beetles physically remove livestock dung from the soil surface, aerate soil, and facilitate dung degradation (Badenhorst et al. 2018; Hajji et al. 2024; Hanafy 2012; Nichols et al. 2008; Tixier et al. 2015). A meta-analysis by Anderson et al. (2024) showed that the presence of dung beetles, on average, increases plant growth by 17% [1%, 35%] (95%CL). In addition, dung beetle activity also reduces the abundance of parasites, such as nematodes and biting flies (deCastro-Arrazola et al. 2023; Nichols et al. 2008). These beneficial effects have long been recognized, leading to the purposeful introduction and release of multiple species, including the bull-horned dung beetle (*Onthophagus taurus*), across the globe (Anderson and Loomis 1978; Pokhrel et al. 2020).

Due to their reliance on vertebrate feces as breeding substrate, dung beetles are vulnerable to residues of veterinary parasiticides and antibiotics in livestock feces, which are known to have both lethal and sublethal effects at ecologically relevant concentrations (e.g.: Hammer et al. 2016; Jacobs and Scholtz 2015; Pérez-Cogollo et al. 2015). Recent studies also reported that livestock dung, including that of cattle, sheep, goats, pigs, and horses, contain various microplastics (Beriot et al. 2021; Sheriff et al. 2023; Wang et al. 2024). However, in contrast to the effects of pharmaceutical residues on dung beetles, the potential consequence of environmental contamination with microplastics on dung beetle survival and behavior remains poorly understood.

Polyurethanes (PU), including thermoplastic polyurethane (TPU), among other plastic types can enter agricultural soils through various points of contamination such as wastewater irrigation, mulch films, and biosolids (Chen et al. 2020; Sahai et al. 2025; Sharmin et al. 2024). Thermoplastic polyurethane, known for its versatility, elasticity, and durability, is ubiquitous in products ranging from footwear and medical devices to industrial films and extrusions (Backes et al. 2024). As part of the broader plastics industry, whose global production surged from approximately 2 million tons in 1950 to over 450 million tons by 2019 (OECD 2022), TPU is a key contributor to the growing demand for synthetic polymers found across diverse sectors. However, standardized methods for quantifying PU/TPU residues in soils remain limited (Yang et al. 2021), highlighting the need for targeted analytical protocols, though some advancements have been made (Choi et al. 2024; Lee et al. 2025). Polyurethane microplastics have been found on pasture soil where concentrations are significantly larger than in grasslands or rangelands (Corradini et al. 2021). This could be due in part to microplastics found in manure (Wang et al. 2024), although the composition and concentration of microplastics in livestock feces is poorly documented.

Environmental contaminants may not only reduce survival directly but also create ‘ecological traps’ (Dwernychuk and Boag 1972) where animals show habitat preference for environments that reduce fitness (Hale and Swearer 2016). For example, some studies show that dung insects prefer ivermectin-contaminated dung over control dung (Errouissi and Lumaret 2010), despite its significant lethal and sublethal effects (Conforti et al. 2018; Jochmann and Blanckenhorn 2016; van Koppenhagen et al. 2020). In dung beetles, ecosystem functions largely depend on maternal substrate choice, tunneling behavior, and investment in each offspring. Because the size of each offspring depends on the size of the brood ball, dung beetles adjust brood ball size and burial depth in response to environmental cues, such as temperature or nutritional quality of the dung (Kirkpatrick and Sheldon 2022; Macagno et al. 2018; Moczek 1998). There is thus significant behavioral plasticity related to the quality of vertebrate dung. Whether females can recognize and avoid microplastics-contaminated dung, or whether contaminated dung acts as an attractant (presenting an ecological trap), is relevant both for larval survival and for the ecosystem services that adults provide.

Here, we use an experimental approach to start investigating whether exposure to thermoplastic polyurethane (TPU) microplastics are a potential concern for dung beetle survival and maternal behaviors. We focus on the widespread dung beetle *Onthophagus taurus*, which has been purposefully introduced for pasture management in large parts of Australia and North America (Anderson and Loomis 1978; Tyndale-Biscoe 1996). Using a laboratory rearing experiment, we first assess how exposure to different concentrations of TPU in cow dung affects larval mortality. Next, we investigate whether adult females can distinguish between or show preferences for contaminated and uncontaminated dung. Although the extent of exposure of dung beetles to TPU is unknown, our study suggests that microplastics may harm dung beetle populations and their ecosystem functioning with consequences for sustainable pasture management and soil health.

## 2. Materials and Methods

### 2.1. Generating TPU microplastics

To begin exploring the impact of microplastics on dung beetles, we tested a single TPU formulation as a representative example, given that TPU microplastics, in general, are likely to be found in agricultural landscapes due to their wide ranging applications. To produce microplastic particles we first melted TPU pellets (BASF Elastollan® #1175A10W) into 76.2 mm × 24.9 mm × 30.0 mm rectangular silicon molds at 175°C. After cooling, these plastic bars were then frozen at −80°C to reduce heating and melting during the grinding process. A bench-mounted belt sander fitted with 40-grit sandpaper was used to grind the plastic bars and produce the resulting microplastics. In accordance with methods used by Allemann et al. (2024), a belt sander was chosen for its widespread availability, affordability, and effectiveness in simulating typical wear abrasion. The resulting microplastics were sifted using a 0.5mm mesh to exclude larger particles.

To quantify the approximate size distribution of the TPU particles, we weighed out 12 different samples ranging in weight from 0.76 mg to 2.64 mg. Samples were placed onto glass slides and photographed using a Pixelink camera (M20C-CYL) attached to a LEICA M205 microscope (darkfield). Calibrated images were processed in ImageJ (Version 1.54m) (Schneider et al., 2012). Images were first converted into binary 8-bit images. To separate touching particles, the watershed function was applied before measurement. Particle size was then quantified with the *Analyze Particles* tool, and Feret’s diameter (the maximum caliper length of each particle) was recorded for all 12 samples independently. The average particle diameter was 0.17 mm with a standard deviation of 0.11 mm. The median was 0.15 mm with a lower and upper quantile of 0.09 mm and 0.23 mm, respectively. The distribution of particle sizes was very similar across all nine independent replicates (see fig. S1).

### 2.2. Effect of TPU microplastic contamination on the survival of dung beetle larvae

We collected *Onthophagus taurus* adults in Santa Ysabel County Preserve to establish laboratory colonies at the University of California, San Diego. Laboratory colonies were maintained under standard conditions and fed once a week with previously frozen cow dung.

To generate individuals used to test the impact of TPU on larval survival, we removed ten mature *O. taurus* females from the laboratory colony and placed them in two standard 5-gallon (~19L) plastic buckets used as ovipositing chambers. Each ovipositing chamber contained a compact sterile mixture of soil and sand and topped with defrosted cow dung (approx. 250g) and were kept in a temperature-controlled insectary at 24°C. After six days, all brood balls were carefully extracted by sifting the soil from the ovipositing chamber. Eggs and larvae were extracted from the maternal brood balls and placed into standardized experimental ‘artificial brood balls’ (as in: Rohner and Moczek 2021; Shafiei et al. 2001). In brief, these artificial brood balls consist of 2 grams (± 0.05 g) of cow dung that were placed in the wells of standard 12-well tissue culture plates (VWR, 734-2324). To simulate natural conditions, we removed excess moisture from the dung by squeezing out liquid through a cheesecloth. All squeezed-out dung was thoroughly homogenized before artificial brood balls were made. 12-well plates containing twelve artificial brood balls were each closed with a lid and frozen until they were used for the experiment.

To simulate contamination with microplastics, we used six different TPU treatments by weight (6% (60 mg g^−1^), 3% (30 mg g^−1^), 1% (10 mg g^−1^), 0.1% (1 mg g^−1^), 0.05% (0.5 mg g^−1^), 0.01% (0.1 mg g^−1^)) as well as a control treatment that did not contain any additional microplastics. These concentrations fall within the range that are commonly used in insect studies (Li et al. 2024). TPU microplastics were weighed out to the appropriate amount using analytical balance (Mettler Toledo, AB135-S) and were then added to artificial brood balls. Each brood ball was thoroughly mixed to ensure that microplastics were well incorporated. It is important to note that we were not able to assess the microplastics concentration of the cow dung that was used to generate the plates.

Estimates of microplastics concentrations in cow dung are poorly studied and reliable field estimates are currently unavailable. Several studies report particles per unit dung, but this is hard to set in context under laboratory conditions. Instead, we use MP concentrations in agricultural or industrial settings as point of reference. The soil samples taken at an industrial site in Australia were found to contain 0.03% - 6.7% of microplastics (Fuller and Gautam 2016). However, compared to other estimates, these values should likely be considered highly contaminated (Büks and Kaupenjohann 2020). In Swiss floodplains, MP concentrations of up to 55.5 mg kg^−1^ were found (Scheurer and Bigalke 2018). Büks and Kaupenjohann (2020) report a global average of 1.7 mg kg^−1^ microplastics in agricultural soils. However, concentrations are often highly variable. While the concentrations used in our experiment are clearly larger than most often encountered in the field, it is very likely that some insects might be exposed to these concentrations.

Eggs were placed individually into treated or control brood balls and incubated at 26 °C. Larval survival was assessed daily. Larvae were considered dead if they showed no movement and no response to gentle probing with featherweight forceps. The experiment was conducted in several blocks. Due to high mortality rates at very high concentrations, not all treatments were represented in every block. To statistically investigate the effect of TPU treatments on larval survival, we used a Cox proportional hazards mixed-effects model fitted via maximum likelihood, as implemented in the R package coxme (Therneau 2022). The TPU treatment was added as categorical fixed effect. Experimental block was included as a random intercept to account for non-independence among individuals reared under the same conditions. Individuals that survived juvenile development and reached the adult stage (i.e., successfully emerged from the pupal cuticle) were treated as censored observations. The model can be expressed as:

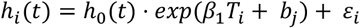

where *h*_*i*_(*t*) is the hazard (i.e., the instantaneous risk of death) of individual *i* at time *t, h*_0_(*t*) is the baseline hazard function, *T* is the effect of TPU treatment. The random effect of block is represented by *b*_*j*_ and the residual error term is indicated with *ε*. The significance of the TPU treatment was assessed via AIC-based model comparison with a null model that included only the intercept and random block effect.

### 2.3. Effect of TPU microplastic contamination on maternal substrate choice

To test the effect of microplastic contamination on maternal behavior, we conducted substrate choice tests. We used standard 5-gallon (~19L) plastic buckets fitted with custom-made wooden dividers to separate the internal volume of the bucket into two equal halves. Autoclaved soil was then added on either side of the divider and compacted until soil was level with the top of the divider. All cow dung was defrosted, drained of excess liquid through a cheesecloth, and homogenized prior to use. To generate microplastic-contaminated dung, 2 ± 0.075 g of sifted TPU microplastics were weighed and added to 200 g of cow dung and thoroughly mixed, resulting in a 1% TPU concentration (1,000 mg kg^−1^). Control portions of approximately 200 g of defrosted cow dung without microplastics were likewise set aside and mixed thoroughly. Both dung treatments were placed on top of the soil on opposite sides of the divider. The two dung pads were separated by about 10 cm to prevent mixing of the two dung types. Because females produce tunnels and brood balls directly underneath the dung, and the two dung sources were separated by the divider, it was possible to distinguish brood balls that were constructed using the contaminated dung from those constructed with control dung on the other side of the divider.

A single *Onthophagus taurus* female was then introduced to the container. To prevent any bias in dung selection, the female was placed in the middle, between the two dung pads. Buckets were covered with mesh to allow for airflow and incubated for six days. Due to logistical constraints, the experiment was repeated in three consecutive temporal blocks. Over the course of three separate experimental blocks, a total of 40 5-gallon buckets were setup with *Onthophagus taurus* females (10 females per block in block 1 and 2, and 20 females in block 3). After six days, soil on either side of the divider was carefully removed and sifted through. Produced brood balls were classified as either made with microplastic or control dung (based on which side of the divider they were collected), weighed (to 2 decimal places), and checked for the presence of an egg or larva. Only brood balls that contained an egg or a larva were used for further analysis to exclude brood balls that were not yet complete.

To test whether females showed a preference for laying brood balls in TPU-spiked or control dung, we fitted a generalized linear mixed-effects model with a binomial error structure. Dung type (TPU vs. control) was treated as a fixed effect, and experimental block was included as a random intercept to account for variation across blocks. To test for an effect on brood ball weight, we fitted log brood ball weight as a function of treatment using mean-centered female size as covariate. Block was again added as a random effect. One brood ball weighing less than 1 g which had large effects on the model was excluded from the analysis.

## 3. Results

### 3.1. Effect of TPU microplastics contamination on the survival of dung beetle larvae

Survivorship of dung beetle larvae declined with increasing concentrations of TPU microplastics (fig. 1). All larvae exposed to TPU concentrations higher than 0.1% died within the first 14 days of their larval stage. Only six percent of all larvae exposed to 0.05% TPU survived to adulthood of (2/36). At the lowest TPU concentration, larval survival was 39% (13/33). In the control treatment, 63% of all larvae survived to adulthood (27/43). Consistent with these proportions, the Cox proportional hazards mixed-effects model indicated a strong effect of the TPU treatment. Including TPU treatment as a fixed effect drastically reduced AIC (ΔAIC = 115.86). Hazard ratios of all treatments were higher than one except for the lowest concentration: The 0.01% TPU treatment had increased mortality, but the hazard ratio was not statistically different from unity (see table 1).

**Table 1:** Hazard ratios (HR) and corresponding 95% confidence limits extracted from a Cox proportional hazards mixed-effects model including experimental block as a random effect.

| treatment | coefficient ($\beta$) | HR [95%CI] | z | P |
| --- | --- | --- | --- | --- |
| 0.01% TPU | 0.44 | 1.56 [0.79, 3.06] | 1.28 | 0.200 |
| 0.05% TPU | 2.33 | 10.28 [5.37, 19.69] | 7.03 | <.001 |
| 0.1% TPU | 2.54 | 12.63 [6.50, 24.55] | 7.48 | <.001 |
| 1% TPU | 3.01 | 20.34 [5.84, 70.85] | 4.73 | <.001 |
| 3% TPU | 3.26 | 25.99 [7.64, 88.42] | 5.21 | <.001 |
| 6% TPU | 2.89 | 18.01 [5.27, 61.63] | 4.61 | <.001 |

**Figure 1:**
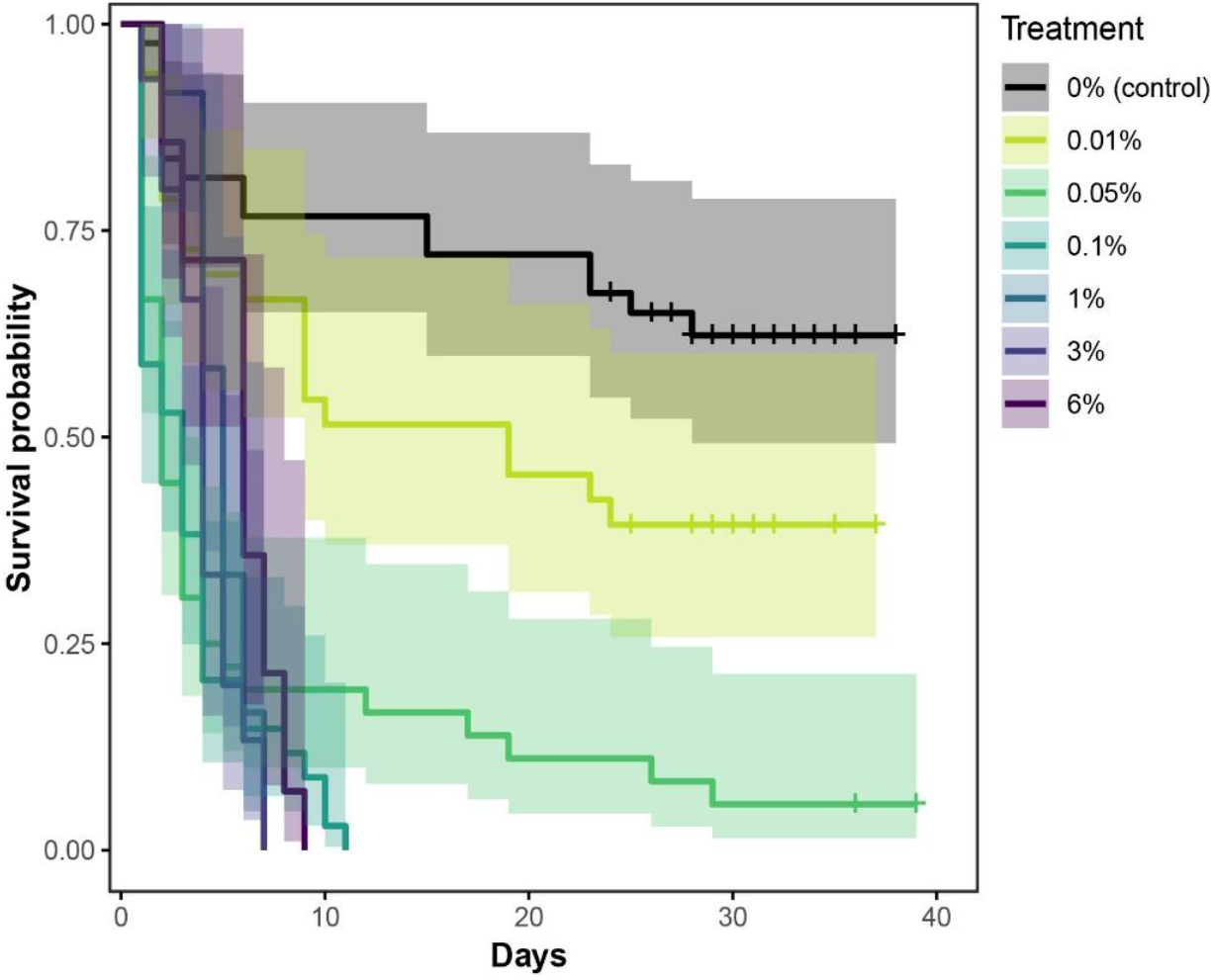
Effect of TPU concentration on larval survival of the dung beetle *Onthophagus taurus*. Vertical tick marks correspond to adult emergence (which were treated as right-censored observations).

### 3.2. Effect of TPU microplastics contamination on maternal substrate choice

During the oviposition choice experiment, 183 brood balls were produced. When provided with the choice between TPU-spiked and control dung, the probability of females using control dung to make brood balls was 0.64 [0.44, 0.79]. More than half of all brood balls were therefore produced with control dung. However, because the 95% confidence limit included 0.5, there was no statistical evidence for significant preference for either source of dung (binomial glmer: z = 1.42, P = 0.157, fig. 2A).

**Figure 2:**
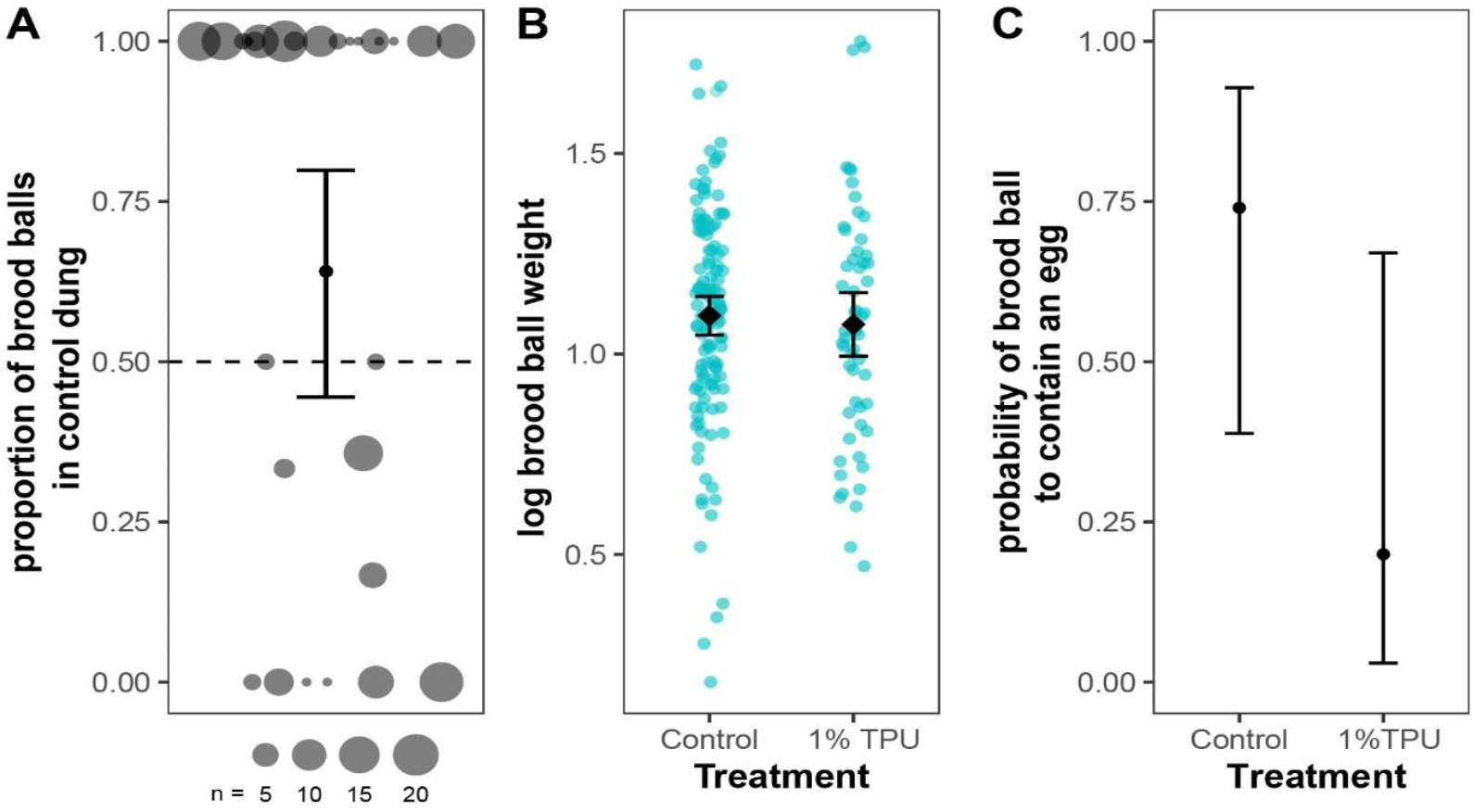
A) Proportion of brood balls produced with control versus TPU-spiked dung in a choice test. Each point represents the proportion of brood balls per female, scaled to the total number of brood balls produced. B) Effect of dung treatment on brood ball size. C) Effect of dung treatment on the likelihood of the brood ball containing an egg. Error bars show 95% confidence limits.

Whether brood balls were produced with control or contaminated dung did not affect their size (lmer: χ^2^_(1)_ = 0.42, P = 0.516, fig. 2B). Adding female body size as a covariate did not change this lack of relationship (χ^2^_(1)_ = 0.09, P = 0.764), although large females produced larger brood balls (χ^2^_(1)_ = 3.92, P = 0.048), as has been shown previously (Kishi 2014). Brood balls produced with control dung overall tended towards a higher likelihood to contain an egg, but this difference was not significant (χ^2^_(1)_ = 1.69, P = 0.194, fig. 2C).

## 4. Discussion

Dung beetles serve as critical detritivores and are routinely exposed to contaminants present in livestock feces. TPU exposure at the concentration range used in our experiment inflicts significant toxic stress that reduces survival of larval dung beetles. We also found no indication that dung beetles behaviorally avoid dung containing microplastics, indicating that their presence poses significant risks to dung beetles and pasture ecosystems. How the concentrations in our experiment compare to exposure in the field remains unknown as reliable studies of microplastics in animal dung have yet to be conducted. However, our results indicate a substantial burden from microplastic exposure that has the potential to impair both dung beetle fitness and their role as ecosystem engineers in agricultural environments.

### 4.1. Potential impact of microplastic pollution on ecosystem functioning

Dung beetles play an important role in agricultural ecosystems. These roles are so important that they have been intentionally introduced to Australia and the United States (amongst other locations) to improve pasture health (Anderson and Loomis 1978; Pokhrel et al. 2021; Tyndale-Biscoe 1996). However, natural populations are now increasingly exposed to chemical stressors and pollutants. Using an experimental approach, we show that exposure to TPU microplastics in cow dung has the potential to drastically reduce larval survival. This potential hazard is exacerbated because reproductively active females do not seem to be able to distinguish between control and TPU-spiked cow dung, similar to how other insects cannot detect chemical pollutants in cow dung (e.g., Conforti et al. 2018). Together, these data suggest that microplastics pollution has the potential to negatively affect dung beetle populations and their ecosystem services. The extent of these impacts, however, is contingent upon the concentration and composition of microplastics present in livestock feces. Several studies report microplastics in livestock dung (Sheriff et al. 2023), but microplastic concentration estimates remain uncertain due to the complexity and high fiber content of samples, reflecting broader methodological challenges with estimating MP concentrations in complex samples. Reported values, often expressed as particles per mass, are difficult to compare to controlled laboratory treatments and may underestimate total loads, especially of smaller particles. Evidence also suggests variation among dung types: for example, cow dung appears relatively low in microplastics compared to dung of other livestock, potentially due to complete digestion during rumination (Zhang et al. 2022b). Whether plastics are truly fully digested or simply become so fragmented that they are undetectable remains unclear. Microplastic abundance in livestock dung is also likely to depend on the type of feed, which often changes over the season. Similar variability has been found in agricultural soils. Comparing different soil samples collected in Denmark, Vollertsen and Hansen (2017) estimated an average of 0.003% microplastics in soil. However, values ranged between 0 and 0.02% (w/w). If similar ranges exist in manure, it is thus very likely that at least some dung beetles will be exposed to significant levels of MPs in the field. Thus, robust field data are required before the impact of microplastics on dung beetle populations can be fully assessed.

Our results may be influenced by the size distribution of microplastic particles (Li et al. 2024). Dung beetles consume a range of particle sizes and microplastics that are too large or too small to be ingested may have less impact. Survival, growth and emergence of aquatic *Chironomus* midges is affected by microplastics, with smaller particulates being more detrimental (Ziajahromi et al. 2018). Our current experiment only tested one particular plastic with one size distribution. Future work on additional types of plastics and size distributions will be needed to assess how varied microplastic sizes impact dung beetles.

Interactions with other stressors will also be important to consider. Chemical pollutants might show complex interactions with other abiotic stressors that should be investigated. Several studies show that microplastics are found in manure that is also contaminated with antibiotics and other pollutants (Mohammadi et al. 2025; Wang et al. 2024). Simultaneous exposure to multiple stressors might cause much stronger interactive (synergistic) effects, as has been found for chemical stressors that interact with temperature (e.g., Gonzalez-Tokman et al. 2022; Walker et al. 2025). Such effects have been found in earthworms where simultaneous exposure to polyethylene MPs and the fungicide carbendazim led to greater reproductive impairment and oxidative‐damage biomarkers than to either stressor alone (Gautam et al. 2024) (see also: Zhang et al. 2020).

### 4.2. Potential mechanisms driving mortality

Irrespective of the environmental exposure, our data suggest that TPU microplastics can drastically reduce survival, but the mechanisms are unclear. Microplastics have been suggested to negatively affect invertebrates through mechanical effects. For instance, in *Drosophila melanogaster* larvae, microplastics can cause physical damage to intestinal cells (Zhang et al. 2020) and similar effects are likely to occur in the non-biting midge *Chironomus riparius* (Prata et al. 2023). Dung beetles might be particularly susceptible to this because they are adapted to feed on hard-to-digest and recalcitrant diets. This includes the development of very large guts, and the repeated ingestion, digestion, excretion, and re-ingestion of the same material. The presence of microplastics could thus damage the gut epithelium and facilitate infection. This is further exacerbated because larvae are physically confined to their brood ball, which limits their ability to feed selectively on potentially contaminated or uncontaminated dung.

An alternative mechanism to physical harm could be chemical toxicity. The TPU we used in this study contains several components that act as plasticizers, chemical additives used to increase flexibility and workability in plastics, including triphenyl phosphate and tricresyl phosphate. Both compounds have been linked to adverse biological effects. For instance, tricresyl phosphate exposure caused intestinal damage and neurotoxicity in earthworms (*Eisenia fetida*) (Yang et al. 2018), while triphenyl phosphate led to DNA damage in the same species (Zhang et al. 2022a). In vertebrates, tricresyl phosphate has been shown to impair fertility in Japanese medaka fish (*Oryzias latipes*) (Chen et al. 2022). The negative effects seen in the present study may thus not only be physical but also caused through chemical leaching. Another potential effect is that microplastics alter the microbiota, as shown in honeybees (Wang et al. 2021) and springtails (Ju et al. 2019). Dung beetles rely on their microbiota for growth, development, and even survival under stressful conditions (Rohner et al. 2024; Schwab et al. 2016). Microplastics-mediated disruption of the microbiome could therefore reduce larval survival. Future work will need to clarify the mechanisms by which microplastics impact arthropod survival, with important implications for environmental management.

Future research should also address the role of dung beetles in the breakdown of microplastics in manure and soil. Because dung beetle larvae use their enlarged mandibles to repeatedly masticate the same material, it is possible that this behavior leads to mechanical fragmentation of the MPs. Similar effects have been found in caddisfly larvae which incorporate plastics into their cases, their case‐building activity can itself fracture larger particles into secondary microplastics (Valentine et al. 2022). In addition, biodegradation mediated by larval gut microorganisms is another possibility in which dung beetles may shape MP. Dung beetles might thus be important mediators of MP contamination and degradation in agricultural landscapes.

## 5. Conclusions

Dung beetles provide important ecosystem services in agricultural landscapes that are increasingly exposed to pollutants that reduce beetle survival and reproduction. We show that TPU microplastic in cow dung significantly reduces survival of dung beetle larvae. Additionally, adult beetles are equally likely to use microplastic-free and contaminated dung, even if contamination levels will expose their offspring to lethal concentrations of microplastics. Because these microplastics are used in agricultural settings and livestock feces are known to contain significant levels of microplastics, it is likely that these negative effects also occur under natural conditions. However, a full environmental hazard assessment will be necessary to fully assess these effects. It will be particularly important to quantify concentrations, composition, size distribution, and variability of microplastics contamination in manure under ecologically relevant conditions.

## Acknowledgements

We are grateful for support by the Department for Ecology, Behavior, and Evolution at the University of California, San Diego.

## Figures

**Figure S1:**
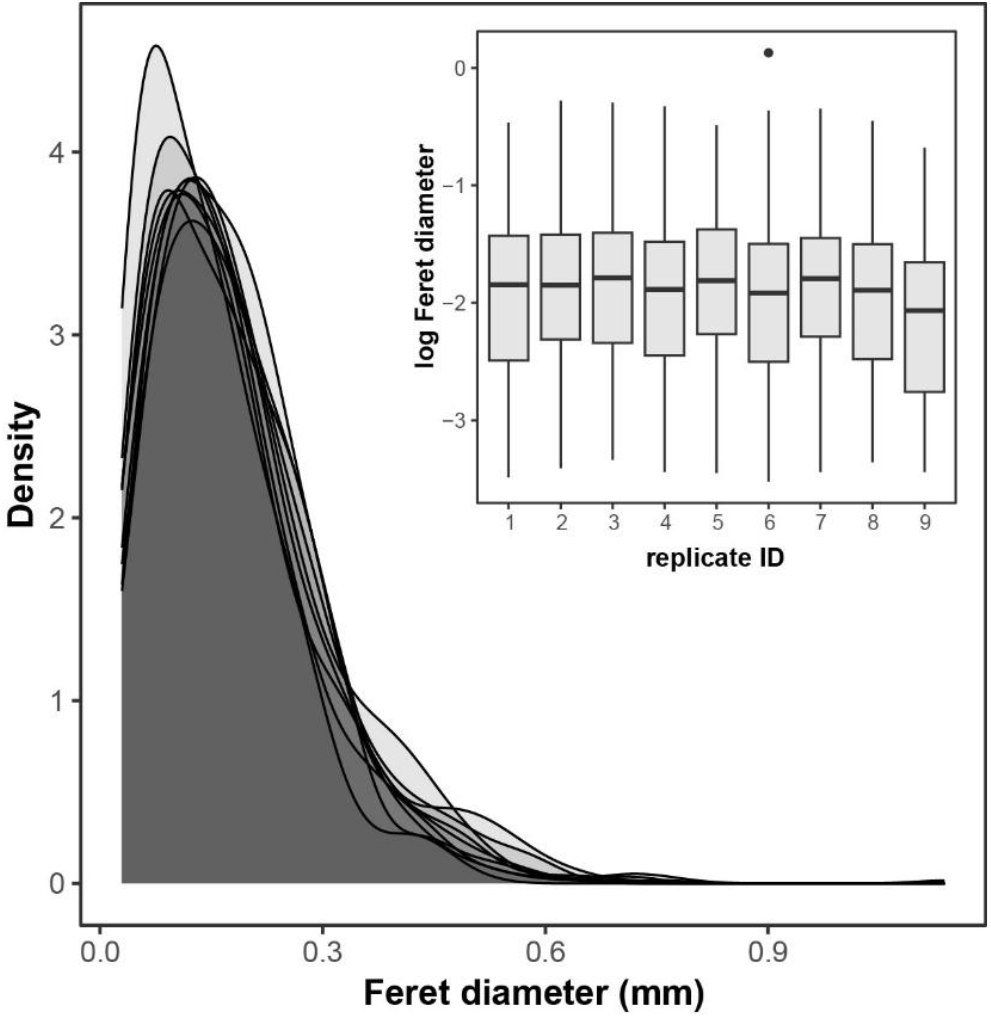
Density distribution of Feret’s diameter (maximum diameter) of TPU microplastics used in this study for 9 independent replicates. Inset shows boxplots (median and quantiles) for all nine replicates on the log scale.

